# Multimodal imaging suggests potential immune-vascular contributions to altered regional brain perfusion and oxygen metabolism in Post-COVID-19 Syndrome

**DOI:** 10.64898/2026.01.12.699018

**Authors:** Daniel Martins, Matthew Burrows, Owen O’ Daily, Ziyuan Cai, Nicole Mariani, Alessandra Borsini, Valeria Mondelli, Brandi Eiff, Silvia Rota, Timothy Nicholson, Laila Rida, Adam Hampshire, Federico Turkheimer, Catherine Morgan, David J. Lythgoe, Steven C.R. Williams, Fernando Zelaya

**Author notes:** **Corresponding author:** Daniel Martins, Department of Neuroimaging, Centre for Neuroimaging Sciences, Institute of Psychiatry, Psychology and Neuroscience, King’s College London, Denmark Hill, SE5 8AF, London, UK.

## Abstract

Post-COVID-19 Syndrome (PCS) frequently presents with persistent fatigue, cognitive impairment, and emotional symptoms. Although structural brain changes remain subtle, growing evidence implicates functional and metabolic disruptions in ongoing symptomatology. We used multimodal MRI to investigate cerebral perfusion and oxygen metabolism in PCS and examined their associations with cognitive function and peripheral biomarkers. We enrolled 40 individuals with prior mild SARS-CoV-2 infection, including 20 with persistent fatigue and 20 recovered controls matched for age, sex, BMI, and acute COVID-19 severity. Participants underwent structural MRI, arterial spin labelling (ASL) to quantify regional cerebral blood flow (CBF), and asymmetric spin echo imaging to estimate oxygen extraction fraction (OEF) and cerebral metabolic rate of oxygen consumption (CMRO□). We assessed cognition using an online battery and measured serum levels of TNF-α, IL-6, IL-8, IL-13, IFN-γ, GFAP, and S100β, alongside blood routine tests. We performed ANCOVAs on predefined regions of interest (hippocampus, anterior cingulate cortex [ACC], insula, amygdala, striatum), followed by Bayesian inference and exploratory whole-brain analyses. PCS participants showed increased CMRO□ in the hippocampus and decreased CMRO□ in the ACC. Subfield analysis revealed elevated OEF and CMRO□ across most hippocampal regions, excluding the entorhinal cortex. Whole-brain analyses identified increased perfusion in salience-related regions (insula, ACC, thalamus) and decreased perfusion in posterior cortical and cerebellar areas, in the absence of grey matter volume differences. Higher hippocampal metabolism positively correlated with cognitive performance, suggesting compensatory adaptation to sustain function. In contrast, lower ACC CMRO□ correlated with depressive symptoms, reduced motivation, and elevated TNF-α and GFAP, consistent with neurovascular uncoupling possibly driven by immune-glial activation. These findings reveal distinct physiological disruptions in PCS, with potential implications for stratified, metabolism-focused interventions.

## Introduction

Post-COVID-19 Syndrome (PCS), or Long COVID, is a heterogeneous and frequently debilitating condition that persists beyond the resolution of acute SARS-CoV-2 infection[1, 2]. Many individuals report enduring symptoms such as fatigue, cognitive impairment, and affective disturbances, which collectively suggest dysfunction in brain systems underlying motivation, memory, attention, and emotion regulation[1–3]. However, the neurobiological mechanisms driving these symptoms remain poorly understood[2, 4, 5].

Structural neuroimaging studies have identified subtle grey matter reductions in PCS patients, particularly in limbic and frontoparietal regions[6–8]. There have been reports of more pronounced alterations in hospitalized vs non-hospitalized patients[9, 10], however alterations have similarly been reported in individuals affected by mild disease only[6] and the extent of these findings remains heterogeneous across cohorts. In contrast, functional neuroimaging has revealed more consistent evidence of disruption. Fluorodeoxyglucose positron emission tomography (FDG-PET) studies frequently show hypometabolism in frontoparietal, medial temporal, and subcortical structures, even in the absence of structural atrophy[11–13], including single-subject sensitivity in several cohorts[14, 15]. These patterns have been interpreted as signatures of neural dysfunction, possibly reflecting immune-mediated metabolic suppression. Likewise, perfusion-based techniques such as arterial spin labelling (ASL) have identified regional hypoperfusion in PCS, particularly affecting frontoparietal regions among individuals with fatigue and cognitive complaints[16].

Although PET studies have provided important insights into hypometabolism in PCS, they primarily reflect glucose uptake, which has been hypothesised to reflect astrocyte-related glutamatergic dysfunction[17, 18]. In contrast, physiological MRI techniques such as ASL and qBOLD offer complementary information by enabling simultaneous, non-radioactive assessment of cerebral blood flow (CBF), oxygen extraction (OEF), and cerebral metabolic rate of oxygen consumption (CMRO[) within the same session[19–21]. These complementary physiological metrics characterize how oxygen is delivered and utilized in the brain, offering insights into potential dissociations between vascular supply and metabolic demand[22]. This integrative approach is particularly well-suited to post-infectious syndromes like PCS, where immune, vascular, and metabolic mechanisms likely converge[5, 23].

In this study, we applied multimodal MRI to quantify resting-state CBF, OEF, and CMRO□ in individuals with PCS and in matched recovered controls (HC). In addition to comparing global and region-specific metrics, we examined whether PCS-related physiological abnormalities were associated with patients’ clinical features, including fatigue severity, cognitive performance, and peripheral markers of inflammation and glial activation.

Based on prior findings of hypoperfusion and hypometabolism in PCS, we hypothesized that affected individuals would show reduced CBF, OEF and CMRO□ in regions supporting cognitive and affective processing. Finally, we anticipated that abnormalities in these physiological measures would correlate with symptom burden, cognitive deviation-from-norm, and circulating immune markers, consistent with a model of neuroimmune dysregulation in PCS.

## Methods

### Study Design and Participants

This study employed a single-site observational case-control design. All participants, both PCS cases and recovered controls, were required to have had biologically confirmed SARS-CoV-2 infection at least three months prior to enrolment, consistent with the WHO clinical case definition of Post-COVID-19 condition[24]. Individuals in the PCS group were characterised by persistent symptoms lasting ≥3 months after infection, whereas recovered controls were required to be fully asymptomatic, with complete resolution of all post-COVID symptoms. We included individuals with mild acute COVID-19 only, to minimise potential confounding effects of hospitalisation, hypoxia, or severe systemic inflammation on brain perfusion and metabolism, and to ensure that any observed group differences were not attributable to the neurological sequelae of moderate–severe disease. All participants were required to have had biologically confirmed SARS-CoV-2 infection prior to enrolment. Confirmation was based on either a positive reverse-transcription polymerase chain reaction (RT-PCR) test from a nasopharyngeal swab or documented serological evidence of prior infection, in accordance with UK national testing guidelines at the time of acute illness. Participants with only clinically suspected but biologically unconfirmed COVID-19 were not included. Recovered healthy control participants were required to have a history of biologically confirmed SARS-CoV-2 infection, followed by complete resolution of all acute and post-acute symptoms. To minimize the risk of including individuals with subtle or subclinical post-COVID manifestations, control participants were screened using the same standardized clinical instruments applied to the PCS group, including the Fatigue Assessment Inventory. Only individuals scoring below established clinical thresholds, and reporting no persistent fatigue, cognitive complaints, affective symptoms, autonomic dysfunction, or post-exertional malaise at the time of assessment, were included as healthy controls. In addition, control participants underwent the same cognitive assessment battery and clinical interview as PCS participants, and individuals demonstrating clinically relevant deviations from normative performance or reporting residual symptoms were excluded. This stringent screening strategy ensured that the control group represented a recovered post-COVID population without ongoing symptomatology, rather than asymptomatic or mildly affected PCS cases.

We recruited participants through King’s College Hospital NHS Trust and surrounding community networks. Eligible individuals were aged 16–65 years, fluent in English, and capable of providing informed consent. We excluded individuals with a history of major neurological or psychiatric disorders, current substance misuse, systemic or central nervous system inflammatory conditions, chronic respiratory or cardiac disease, recent vaccination (within two weeks), pregnancy or breastfeeding, BMI >30, or current use of immunomodulatory or psychoactive medications. We used the global severity score of Fatigue Assessment Inventory (FAI) to pre-screen participants for symptom severity [25]. Individuals scoring greater than 4 on the global fatigue severity scale were classified into the PCS group with persistent fatigue, while those scoring 4 or below were included as recovered controls. The threshold of > 4 was selected to ensure inclusion of individuals with clinically meaningful levels of fatigue, consistent with prior applications of the FAI in post-viral and chronic fatigue contexts[25]. Groups were balanced for age, sex, BMI, and acute COVID-19 severity. The study received ethical approval from the UK Health Research Authority (IRAS ID: 308661) and was conducted in accordance with the Declaration of Helsinki and Good Clinical Practice (GCP) guidelines. All participants provided written informed consent prior to study procedures.

We conducted an a priori power analysis using G*Power (version 3.1) to estimate the required sample size for detecting group differences in dynamic functional connectivity metrics using analysis of covariance (ANCOVA). Assuming a medium to large effect size (Cohen’s f=0.30, equivalent to ηp^2^≈0.08), alpha = 0.05, power (1 − β) = 0.80, and three covariates (age, sex, and handedness), the estimated sample size required per group was 19. Our sample of 20 participants per group therefore provides adequate power to detect effects in this range.

### Procedures

Each participant completed a single study visit that included clinical assessments, venous blood collection, and a multimodal MRI scan. Clinical assessments included medical and psychiatric history, physical examination, vital signs, and a structured battery of standardized questionnaires. Fatigue was assessed using both the full Fatigue Assessment Inventory[25] and the Multidimensional Fatigue Inventory[26] to differentiate between mental and physical fatigue. Symptoms of autonomic dysfunction were evaluated using the Composite Autonomic Symptom Score (COMPASS-31)[27], while mood and anxiety were measured using the Hospital Anxiety and Depression Scale (HADS)[28]. Sleep quality was assessed with the Pittsburgh Sleep Quality Index[29], and respiratory symptoms were rated using the MRC Dyspnoea Scale[30]. Additional measures included the DePaul Symptom Questionnaire[31], the American College of Rheumatology (ACR) fibromyalgia criteria[32], the PAIN Detect questionnaire[33], and Montreal Cognitive Assessment[34]. Most self-report questionnaires were completed remotely via the Qualtrics platform within 72 hours of the in-person visit. All participants underwent a structured diagnostic assessment using the Mini International Neuropsychiatric Interview (MINI) to exclude current or past major psychiatric disorders, including mood, anxiety, psychotic, and substance use disorders.

### Neuroimaging data acquisition and processing

Neuroimaging data were acquired on a 3T GE Healthcare MR750 scanner (GE Medical Systems, Milwaukee, WI), using a 32-channel receive-only head coil. The protocol included high-resolution anatomical imaging, cerebral perfusion quantification, and quantitative BOLD imaging to assess tissue oxygenation and metabolism. Additional MRI sequences were acquired as part of the broader study’s imaging protocol and will be reported elsewhere.

Structural images were obtained using a 3D T1-weighted magnetisation-prepared rapid gradient echo (MPRAGE) sequence with the following parameters: repetition time (TR) = 2300 ms, echo time (TE) = 2.98 ms, inversion time (TI) = 900 ms, and isotropic voxel size of 1 mm³. These images were used for anatomical reference, spatial normalisation and also sensitivity analyses exploring whether metabolic differences between groups could reflect changes in grey matter volume. For the later, structural images were preprocessed using SPM12 and the Computational Anatomy Toolbox (CAT12)[35]. Images were segmented into grey matter, white matter, and cerebrospinal fluid and normalized to MNI space using DARTEL[36]. Grey matter volume was computed using voxel-based morphometry[37].

Regional CBF was quantified using a 3D pseudo-continuous arterial spin labelling (pCASL) sequence with a labelling duration of 1.8 seconds and a post-labelling delay of 2.0 seconds. Background suppression and a segmented 3D fast spin echo stack-of-spirals readout were employed to enhance signal-to-noise ratio and reduce motion artefacts. A matched proton density scan was also acquired to enable absolute quantification of CBF. Acquisition time for the 3D pCASL sequence was 6:15min.

Voxelwise estimates of OEF from the transverse relaxation rate sensitive to deoxygenated blood volume (R2′) were obtained using a multi-echo asymmetric spin echo (ASE) sequence as part of a qBOLD modelling framework, following the methodology proposed by Yablonski et al[38]; and Stone et al[39]. The sequence employed 17 shifts of the 180deg pulse starting from 0 to 32ms in steps of 2ms. Gradient shimming along the z-axis was incorporated into the sequence to correct for macroscopic magnetic field inhomogeneities. Nine ‘z-shimming’ gradients (-100 to +100 uT/m in steps of 25uT/m] were applied before each echo shift of the ASE[40], to allow OEF estimation from computation of R2′. Acquisition time for the ASE sequence was 8:25min. Both CBF and OEF maps were normalized to MNI space without spatial smoothing in order to preserve voxel-level precision for metabolic calculations. Because CBF images were processed using the DARTEL framework, which retains native voxel resolution (1.88 × 1.88 × 3 mm), the OEF maps - initially skull-stripped using resliced intracranial volume (ICV) masks - were resampled to match the CBF dimensions to ensure voxelwise correspondence.

The cerebral metabolic rate of oxygen consumption (CMRO[) was computed using the Fick principle[41] and expressed in units of μmol O[/100 g tissue/min. For each participant, CMRO□ was calculated at the voxel level as the product of CBF, OEF, and arterial oxygen content (CaO[). The CBF and OEF values were extracted from the aligned and normalised maps, while CaO□ was derived using subject-specific haemoglobin concentration (Hb, reported in g/100 mL) determined from venous blood samples according to the formula: CaO□ = [ (Hb × 1.34 × SaO□) + (0.0031 × pO□) ] × 44.64 / 100. In this equation, SaO□ was assumed to be 0.98, pO□ was fixed at 100 mmHg, 1.34 mL O[/g is the oxygen-carrying capacity of haemoglobin, and 44.64 is the molar conversion factor from millilitres of O□ to micromoles. Voxelwise multiplication of CBF and OEF maps was performed using a custom MATLAB script, followed by scaling with the subject-specific CaO□ value using a second script.

All maps were co-registered to native T1 space, normalized to MNI space, and smoothed using a 6 mm FWHM Gaussian kernel. Global physiological values were extracted using a grey matter probability mask (P(grey matter)>0.2), while regional values were obtained from bilateral ROIS defined based on the Harvard-Oxford atlas, including the hippocampus, anterior cingulate cortex (ACC), amygdala, and striatum[42], and on the Hammersmith atlas for the anterior and posterior insula[43]. We decided to include bilateral ROIs instead of lateralised for two main reasons: first, we had no reason to suspect of lateralised effects; second, by averaging across a higher number of voxels we would maximise SNR. We also investigated hippocampal subfields as previous evidence have suggested subregional structural and functional alterations in PCS[44, 45]. For that purpose, we derived subject specific masks using the hippocampal subfield segmentation pipeline from FreeSurfer 7.1[46], merging smaller regions to create functionally interpretable masks corresponding to the Cornus Ammonis (CA1-3), dentate gyrus (including CA4), subiculum (including presubiculum and parasubiculum) and entorhinal cortex (including granule and molecular layers). ROI selection was hypothesis-driven and grounded in existing PCS neuroimaging literature. FDG-PET studies of PCS have repeatedly reported hypometabolism in the anterior cingulate cortex, insula, medial temporal lobe, striatum, and frontoparietal regions[11, 12]. ASL-MRI studies have similarly identified hypoperfusion affecting frontoparietal and salience-network regions among individuals with fatigue and cognitive complains[16, 47]. These regions also align closely with the core symptom domains of PCS, including episodic memory dysfunction (hippocampus), motivational and affective disturbances (ACC), interoceptive and autonomic symptoms (insula), and mood-related or stress-sensitivity features (amygdala and striatum)[48]. In addition, recent high-resolution MRI studies have described hippocampal subfield alterations in PCS and other post-viral fatigue syndromes[44, 45], which motivated our inclusion of subfield-specific analyses.

Quality control procedures were implemented at multiple stages to ensure anatomical alignment, physiological plausibility, and robustness of the derived quantitative maps. All CBF, OEF, and CMRO□ images were visually inspected following coregistration to confirm accurate alignment to structural T1-weighted images and to each other. Special care was taken to verify that OEF maps, which were resampled to match the native resolution of the CBF images following DARTEL processing, retained spatial correspondence across modalities. In addition to visual inspection, extracted values were reviewed to confirm biological plausibility; values falling outside established physiological ranges were flagged and excluded where necessary. To maintain the integrity of spatial specificity, median values within each ROI were extracted from unsmoothed CBF, OEF, and CMRO□ maps using the fslstats tool. For each ROI, we assessed signal quality and completeness by computing the proportion of voxels with values below a conservative physiological threshold for OEF (<0.05), and ROIs with more than 75% of such voxels were excluded from subsequent statistical analyses. These procedures ensured that all reported regional metrics were based on anatomically accurate, physiologically valid, and quantitatively reliable data.

### Blood analysis

Venous blood samples (up to 30 mL) were collected on the day of imaging. Standard hematological and biochemical panels - including erythrocyte sedimentation rate (ESR), C-reactive protein (CRP), thyroid-stimulating hormone (TSH), free thyroxine (T4), and liver function tests - were processed by Synnovis (Viapath) at King’s College Hospital NHS Foundation Trust. Serum samples were frozen at −80°C and later assayed for cytokines and glial markers using electrochemiluminescence immunoassays (Meso Scale Discovery, Rockville, MD, USA). The cytokine panel included interleukin-1 beta (IL-1β), interleukin-2 (IL-2), interleukin-4 (IL-4), interleukin-6 (IL-6), interleukin-8 (IL-8), interleukin-10 (IL-10), interleukin-12p70 (IL-12p70), interleukin-13 (IL-13), tumor necrosis factor-alpha (TNF-α), and interferon-gamma (IFN-γ). Glial markers included glial fibrillary acidic protein (GFAP), measured with the ultrasensitive kit, and S100β. Assays were performed in duplicate. Values below the detection limit were discarded. This led us to discard all quantifications of IL-2, IL-4 and IL-12p70. Outliers greater than three standard deviations from the group mean were winsorized. No analyte had more than 10% missing data, therefore all data were included.

### Cognitron

Cognitive performance was assessed online using a battery of computerized tasks from the Cognitron platform, covering delayed verbal memory, working memory, executive function, attention, motor coordination, and spatial manipulation. Tasks included Verbal Analogies (abstract reasoning and semantic integration), Immediate and Delayed Prospective Memory (episodic memory), 2D Spatial Manipulation (visuospatial working memory and mental rotation), Motor Control (sensorimotor coordination and reaction consistency), Spotter (vigilance and sustained attention), Lead Balloon (abstract reasoning and effort), and Block Reasoning (fluid intelligence and rule-based problem solving). Each task involves trial-level response collection, capturing both accuracy and reaction time. Performance metrics were transformed into standardized deviation-from-expected (DFE) scores using normative models derived from a large independent sample (n > 20,000), adjusted for age, sex, handedness, and harmonized ethnicity. DFE scores represent standardized residuals (observed minus expected performance divided by normative standard deviation) and were computed separately for accuracy and reaction time.

### Statistical analysis

Statistical analyses were conducted using JAMOVI (version 2.3) and SPM12. Between-group comparisons of global and regional physiological measures, including CBF, OEF, and CMRO[, were performed using general linear models, with age, sex, and handedness included as covariates. Bayesian hypothesis testing was conducted in parallel to quantify the relative strength of evidence for the null versus the alternative hypothesis. Bayes factors (BF□□) were used to interpret the results, where values greater than 1 indicate support for the alternative hypothesis, and values less than 1 support the null. Specifically, BF□□ values between 1 and 3 were considered to provide anecdotal evidence for the alternative, between 3 and 10 as moderate evidence, and above 10 as strong evidence. Conversely, BF□□ values between 0.33 and 1 were considered anecdotal evidence for the null, between 0.1 and 0.33 as moderate, and below 0.1 as strong evidence for the null hypothesis. This Bayesian framework allowed us to distinguish true null effects from inconclusive findings due to limited power.

Whole-brain group comparisons on smoothed parametric maps were performed using two-sample t-tests in SPM12, with a cluster-forming threshold set at p < 0.001 (uncorrected) and family-wise error (FWE) correction applied at the cluster level with pFWE < 0.05. These analyses encompassed the entire brain, including cortical, subcortical, cerebellar, and brainstem grey matter, as defined by the unified SPM12 tissue segmentation pipeline and the corresponding grey matter probability maps. Our aim was to test for any additional group differences beyond those identified a priori in the ROI analysis, enabling the detection of exploratory perfusion or metabolic alterations across the whole brain. To investigate whether perfusion or metabolic group differences could reflect neurodegenerative alterations, we performed sensitivity analyses on estimated regional grey matter volume while accounting for total intracranial volume (TIV) as additional covariate. Associations between imaging-derived physiological measures, cognitive deviation-from-expected (DFE) scores, clinical symptom scales, and serum biomarkers were examined using partial Spearman rank correlations, adjusting for age, sex, handedness, group and global CBF/OEF/CMRO_2_ as appropriate. Given the exploratory nature of these analyses, correlation results are reported without correction for multiple comparisons and should be interpreted with caution.

## Results

### Sociodemographics and clinical characterisation

The groups were matched for age, sex, ethnicity, and body mass index (BMI), with no significant differences observed across these variables (all p > 0.5; Table 1). However, groups differed significantly in years of education, with HC participants reporting higher educational attainment. PCS participants also had a significantly longer duration since first SARS-CoV-2 infection, and were significantly less likely to have been vaccinated prior to infection. Clinically, individuals with PCS reported significantly greater symptom burden across multiple domains. These included fatigue (both visual analogue scale and multidimensional components), sleep disturbance (PSQI), post-exertional malaise, autonomic dysfunction (COMPASS-31), affective symptoms (HADS depression/anxiety), PTSD-related symptoms, and musculoskeletal complaints (WPI and SS scores). Notably, 40% of PCS participants met criteria for fibromyalgia, compared to none in the HC group. These differences were supported by both frequentist (p < 0.05) and Bayesian inference, with many outcomes yielding strong to extreme evidence in favor of group separation (BF□□ > 10), particularly for fatigue, post-exertional malaise, and pain sensitivity (Supplementary Table S1).

**Table 1.**
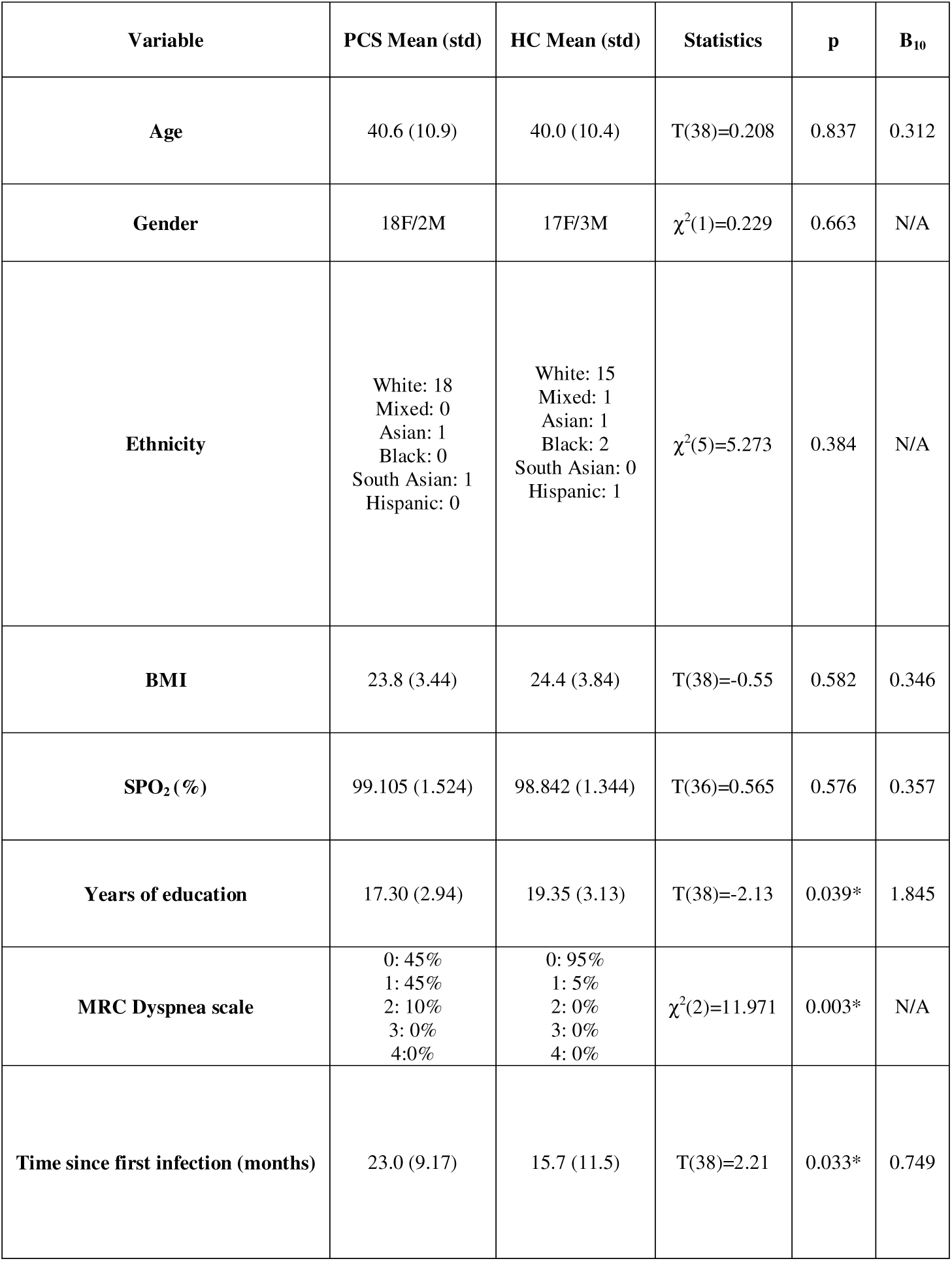
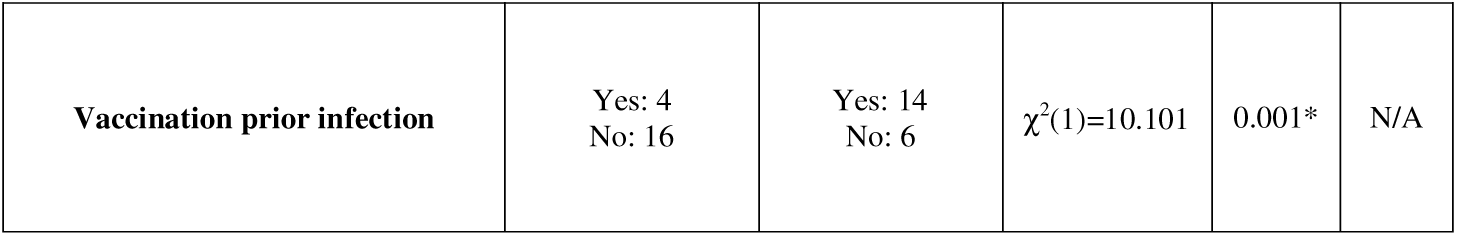
Sociodemographic and clinical characteristics of PCS and healthy control participants. Group comparisons of demographic and COVID-19-related variables between individuals with Post-COVID-19 Syndrome (PCS) and matched healthy controls (HC). Data are shown as mean (standard deviation) or counts. Between-group differences were assessed using independent samples t-tests or chi-squared tests as appropriate. Bayes factors (BF□□) quantify evidence for group differences; values >1 indicate increasing support for the alternative hypothesis. Asterisks (*) denote statistically significant results at p < 0.05 or BF□□ > 1.

| Variable | PCS Mean (std) | HC Mean (std) | Statistics | p | B<sub>10</sub> |
| --- | --- | --- | --- | --- | --- |
| Age | 40.6 (10.9) | 40.0 (10.4) | T(38)=0.208 | 0.837 | 0.312 |
| Gender | 18F/2M | 17F/3M | $\chi^2(1)=0.229$ | 0.663 | N/A |
| Ethnicity | White: 18<br>Mixed: 0<br>Asian: 1<br>Black: 0<br>South Asian: 1<br>Hispanic: 0 | White: 15<br>Mixed: 1<br>Asian: 1<br>Black: 2<br>South Asian: 0<br>Hispanic: 1 | $\chi^2(5)=5.273$ | 0.384 | N/A |
| BMI | 23.8 (3.44) | 24.4 (3.84) | T(38)=-0.55 | 0.582 | 0.346 |
| SPO<sub>2</sub> (%) | 99.105 (1.524) | 98.842 (1.344) | T(36)=0.565 | 0.576 | 0.357 |
| Years of education | 17.30 (2.94) | 19.35 (3.13) | T(38)=-2.13 | 0.039* | 1.845 |
| MRC Dyspnea scale | 0: 45%<br>1: 45%<br>2: 10%<br>3: 0%<br>4: 0% | 0: 95%<br>1: 5%<br>2: 0%<br>3: 0%<br>4: 0% | $\chi^2(2)=11.971$ | 0.003* | N/A |
| Time since first infection (months) | 23.0 (9.17) | 15.7 (11.5) | T(38)=2.21 | 0.033* | 0.749 |
| **Vaccination prior infection** | Yes: 4<br>No: 16 | Yes: 14<br>No: 6 | $\chi^2(1)=10.101$ | 0.001* | N/A |

### Clinical routine blood tests

Group comparisons of routine blood parameters revealed no major differences between PCS and HC participants (Supplementary Table S2). However, PCS participants showed numerically higher creatinine and Gamma-GT levels, with corresponding Bayes factors (BF□□ = 1.232 and 1.735, respectively) providing anecdotal to moderate evidence for group differences. Platelet count also trended higher in the PCS group (p = 0.08; BF□□ = 1.123). Other metabolic, inflammatory, and hematological markers - including C-reactive protein, white and red cell counts, hemoglobin, and thyroid function - did not differ significantly between groups.

### Serum cytokines and glial markers

No statistically significant differences were observed in circulating levels of IFN-γ, IL-1β, IL-6, IL-8, IL-10, IL-13, TNF-α or GFAP (Supplementary Table S3). However, TNF-α (p = 0.116; BF□□ = 1.094) and S100β (p = 0.149; BF□□ = 1.025) showed suggestive evidence of elevation in the PCS group, consistent with mild glial or immune activation.

### Cognitive performance

Cognitive testing using the Cognitron battery revealed subtle but functionally relevant differences between groups (Supplementary Table S4). PCS participants showed significantly poorer performance on the delayed object memory task (RT_DFE; p = 0.02; BF□□ = 2.653), as well as trend-level impairments in the Lead Balloon task (abstract reasoning; p = 0.06; BF□□ = 1.330) and vigilance (Spotter task RT; p = 0.06; BF□□ = 1.303). Other domains - including motor control, 2D spatial manipulation, and verbal analogies - did not show significant differences. Overall, Bayesian analysis provided moderate evidence for reduced memory and attentional function in PCS.

### Global CBF, OEF and CMRO_2_

Group-level comparison of median global CBF, OEF and CMRO_2_ extracted from a grey-matter mask revealed no significant difference between individuals with PCS and healthy recovered controls for the three measures, after adjusting for age, gender, and handedness. The absence of group differences was generally supported by our bayesian tests, which showed that the null hypothesis was more likely than the alternative hypothesis given the data (Table 2). Similarly, we could not detect any evidence of qualitative abnormalities in the respective maps of any subject.

**Table 2.** Global cerebral perfusion and oxygen metabolism in PCS and healthy control participants. Group-level comparisons of global grey matter cerebral blood flow (CBF), oxygen extraction fraction (OEF), and cerebral metabolic rate of oxygen consumption (CMRO□ between individuals with Post-COVID-19 Syndrome (PCS) and healthy controls (HC). Values reflect results from ANCOVA models adjusting for age, sex, and handedness. Bayes Factors (BF□□) quantify the strength of evidence for group differences; values <1 support the null hypothesis. No significant group differences were observed at the global level.

| Variable | Statistics | BF<sub>10</sub> |
| --- | --- | --- |
| Global CBF | F(1,35)=2.080, p=0.158 | 0.748 |
| Global OEF | F(1,35)=0.021, p=0.887 | 0.334 |
| Global CMRO<sub>2</sub> | F(1,35)=0.628, p=0.433 | 0.405 |

**Figure 1.**
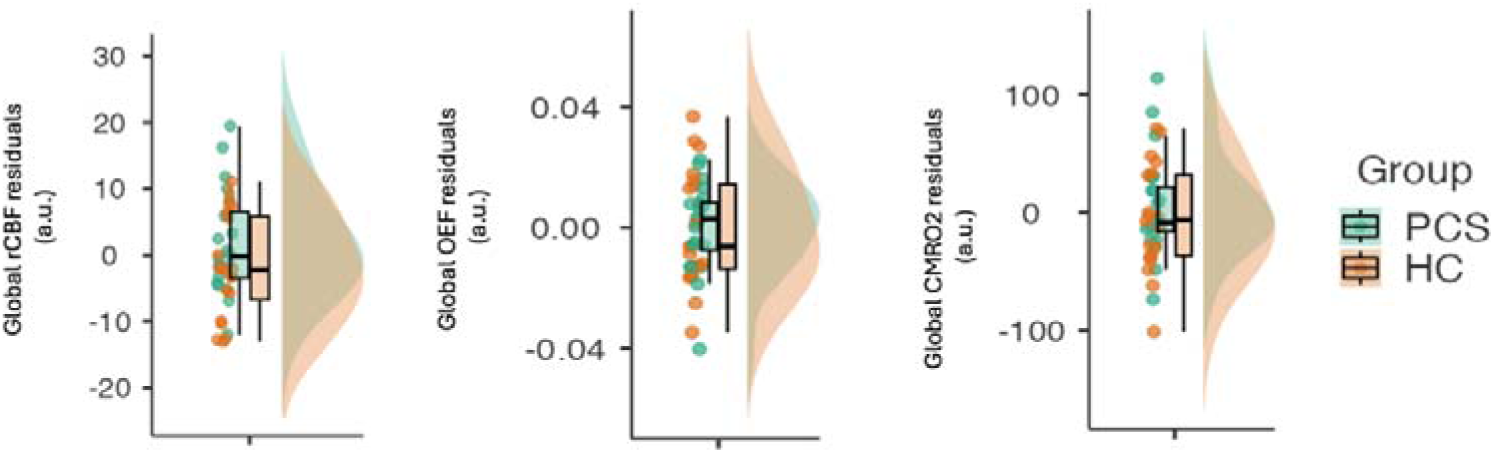
Global estimates of cerebral perfusion and oxygen metabolism in Post-COVID-19 syndrome (PCS). Group-level comparisons of median global cerebral blood flow (CBF), oxygen extraction fraction (OEF), and cerebral metabolic rate of oxygen consumption (CMROL) showed no significant differences between individuals with PCS and recovered healthy controls. These comparisons were adjusted for age, gender, and manual dexterity. Thus, plots show the residuals of the corresponding variables after the effects of co-variate of non-interest were removed. Bayesian analysis similarly supported the null hypothesis, suggesting no strong evidence for group-level differences at the global level.

### Regional CBF, OEF and CMRO_2_ (hypothesis-driven analysis)

For rCBF and OEF, we could not find any significant differences between the two groups for any of our chosen bilateral ROIs (amygdala, striatum, ACC, anterior and posterior insula, or hippocampus). However, for CMRO_2_, we found significant decreases and increases in PCS compared to recovered controls for the ACC and hippocampus ROIs, respectively (Table 3).

**Table 3.** Regional CBF, OEF, and CMRO□ in predefined functionally relevant brain areas. Group comparisons of regional cerebral blood flow (rCBF), oxygen extraction fraction (OEF), and cerebral metabolic rate of oxygen consumption (CMRO[) in bilateral regions of interest (ROIs): amygdala, rostral anterior cingulate cortex (ACC), insula (anterior and posterior), striatum, and hippocampus. ANCOVA models included age, sex, and handedness as covariates. Bayes Factors (BF□□) quantify the strength of evidence in support of group differences. Statistically significant results (p < 0.05 or BF□□ > 1) are marked with an asterisk (*). PCS participants exhibited significantly reduced CMRO□ in the rostral ACC and elevated CMRO□ in the hippocampus, with corresponding moderate-to-strong Bayesian evidence. Trends were also observed in hippocampal and ACC OEF.

| Metric | Region | Statistics | BF<sub>10</sub> |
| --- | --- | --- | --- |
| rCBF | Amygdala | F(1,35)=0.033, p=0.856 | 0.313 |
| | Rostral cingulate | $F(1,35)=0.001, p=0.992$ | 0.314 |
| | Anterior Insula | $F(1,35)=0.088, p=0.768$ | 0.330 |
| | Posterior insula | $F(1,35)=0.021, p=0.885$ | 0.320 |
| | Striatum | $F(1,35)=0.010, p=0.921$ | 0.319 |
| | Hippocampus | $F(1,35)=0.117, p=0.735$ | 0.329 |
| OEF | Amygdala | $F(1,34)=2.524, p=0.121$ | 0.905 |
| | Rostral cingulate | $F(1,35)=3.933, p=0.055$ | 1.607* |
| | Anterior Insula | $F(1,35)=0.407, p=0.528$ | 0.386 |
| | Posterior insula | $F(1,35)=0.591, p=0.447$ | 0.394 |
| | Striatum | $F(1,35)=0.337, p=0.565$ | 0.359 |
| | Hippocampus | $F(1,35)=2.975, p=0.094$ | 1.030* |
| CMRO2 | Amygdala | $F(1,34)=1.914, p=0.176$ | 0.622 |
| | Rostral cingulate | $F(1,35)=4.776, p=0.036^*$ | 1.587* |
| | Anterior Insula | $F(1,35)=0.324, p=0.573$ | 0.378 |
| | Posterior insula | $F(1,35)=0.589, p=0.448$ | 0.372 |
| | Striatum | $F(1,35)=0.336, p=0.851$ | 0.313 |
| | Hippocampus | $F(1,35)=6.927, p=0.013^*$ | 5.056* |

**Figure 2.**
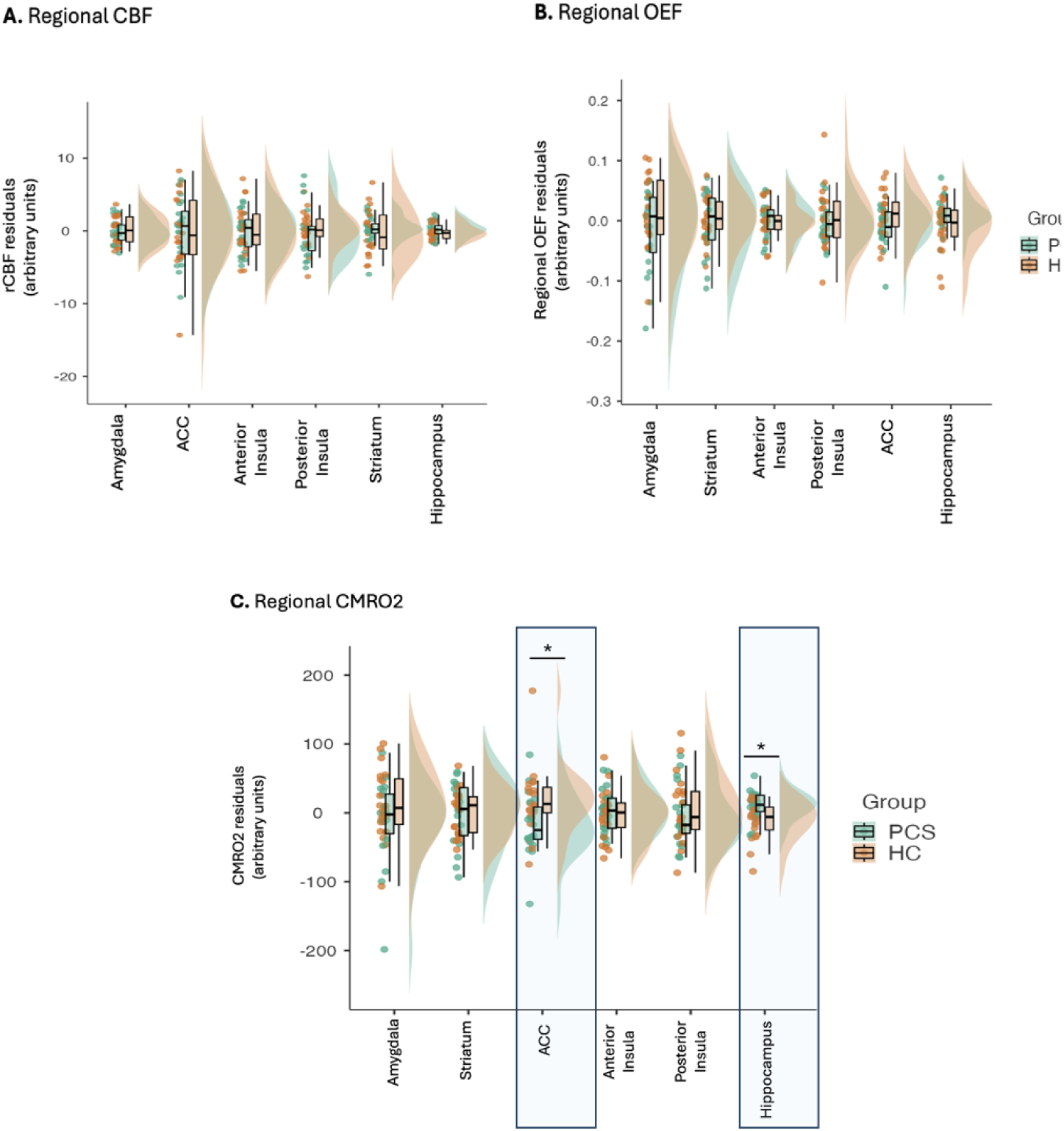
Region-of-interest comparisons of cerebral perfusion and oxygen metabolism in functionally relevant brain regions. Median regional estimates of CBF, OEF, and CMROL were extracted from bilateral anterior cingulate cortex (ACC), amygdala, striatum, hippocampus, and anterior and posterior insula regions-of-interest. Analyses were adjusted for age, gender, dexterity and global CBF, OEF or CMRO_2_ as appropriate. Therefore, here we show the residuals of the corresponding variables after the effects of the covariates of non-interest were removed. While CBF and OEF showed no significant group differences, CMROL was significantly altered in PCS compared to controls, with increases observed in the hippocampus and decreases in the ACC. * p<0.05.

Within the hippocampus, we also inspected four sub-regions: cornus ammonis, dentate gyrus, entorhinal cortex and subiculum. For rCBF, we could not find group differences for any of the hippocampal subregions ROIs. For OEF and CMRO_2_, we found significant increases in PCS compared to recovered controls for all hippocampal subregions, with the exception of the entorhinal cortex, where we could not find any significant group difference for either metric. The results of these frequentist analyses were largely supported by the corresponding bayesian counterparts (Table 4).

**Table 4.** Subregional CBF, OEF, and CMRO□ estimates across hippocampal subfields in PCS and HC participants. Group comparisons of cerebral blood flow (rCBF), oxygen extraction fraction (OEF), and cerebral metabolic rate of oxygen consumption (CMRO[) across four bilateral hippocampal subfields: Cornu Ammonis (CA1–3), Dentate Gyrus (DG), Entorhinal Cortex (EC), and Subiculum. Statistical comparisons were adjusted for age, sex, handedness and global CBF, OEF or CMRO_2_ as appropriate. Bayes Factors (BF□□) reflect the strength of evidence in support of group differences, with values >1 indicating support for the alternative hypothesis. PCS participants showed significantly increased OEF and CMRO□ in CA, DG, and Subiculum, but not in the entorhinal cortex. These results support the presence of subfield-specific metabolic upregulation in PCS. Asterisks (*) indicate statistical significance at p < 0.05 or BF□□ > 1.

| Metric | Region | Statistics | BF<sub>10</sub> |
| --- | --- | --- | --- |
| rCBF | Cornu Ammonis | F(1,35)=0.107, p=0.746 | 0.338 |
|  | Dentate gyrus | F(1,35)=0.253, p=0.618 | 0.350 |
|  | Entorhinal Cortex | F(1,35)=0.009, p=0.927 | 0.315 |
|  | Subiculum | F(1,35)=0.185, p=0.670 | 0.335 |
| OEF | Cornu Ammonis | F(1,35)=4.159, p=0.049* | 1.733* |
|  | Dentate gyrus | F(1,35)=4.700, p=0.037* | 2.203* |
|  | Entorhinal Cortex | F(1,35)=0.307, p=0.583 | 0.360 |
|  | Subiculum | F(1,35)=4.337, p=0.045* | 1.873* |
| CMRO<sub>2</sub> | Cornu Ammonis | F(1,35)=5.921, p=0.020* | 3.408* |
|  | Dentate gyrus | F(1,35)=5.647, p=0.023* | 3.115* |
|  | Entorhinal Cortex | F(1,35)=0.266, p=0.609 | 0.335 |
|  | Subiculum | F(1,35)=9.553, p=0.004* | 12.405* |

**Figure 3.**
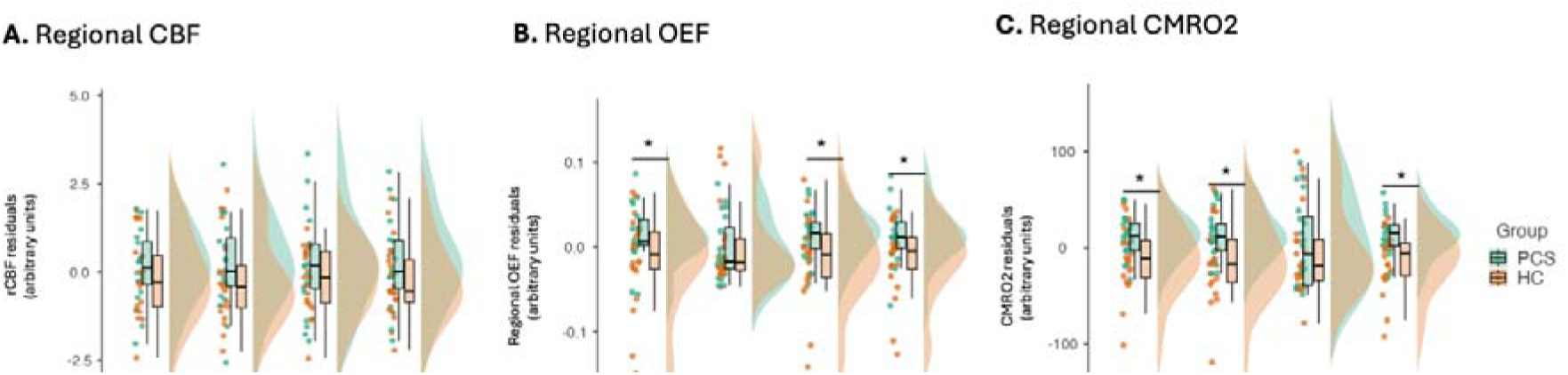
Cerebral perfusion and oxygen metabolism subregional analysis of the hippocampus. Analysis of four hippocampal subfields - cornus ammonis, dentate gyrus, entorhinal cortex, and subiculum - revealed significantly increased OEF and CMROL in PCS compared to controls in all regions except the entorhinal cortex. Here we show the residuals of the corresponding variables after the effects of the covariates of non-interest were removed. * p<0.05.

### Whole-brain rCBF, OEF and CMRO_2_ (exploratory analysis)

For rCBF, we found four significant clusters where PCS showed higher rCBF than controls. These clusters were located primarily in the middle and anterior cingulate cortex (cluster 1), right insula, superior temporal and precentral gyri (cluster 2), left insula, thalamus and caudate (cluster 3), and right middle and inferior frontal gyri, right insula and putamen (cluster 4). We also found two clusters where PCS showed lower rCBF than controls. These clusters spanned the left posterior cingulate, occipital gyri, cerebellum (cluster 5) and left parahippocampal gyrus and striatum (cluster 6) (Supplementary Table S5). For the OEF and CMRO_2_, no clusters survived correction for multiple comparison at the whole-brain level.

**Figure 4.**
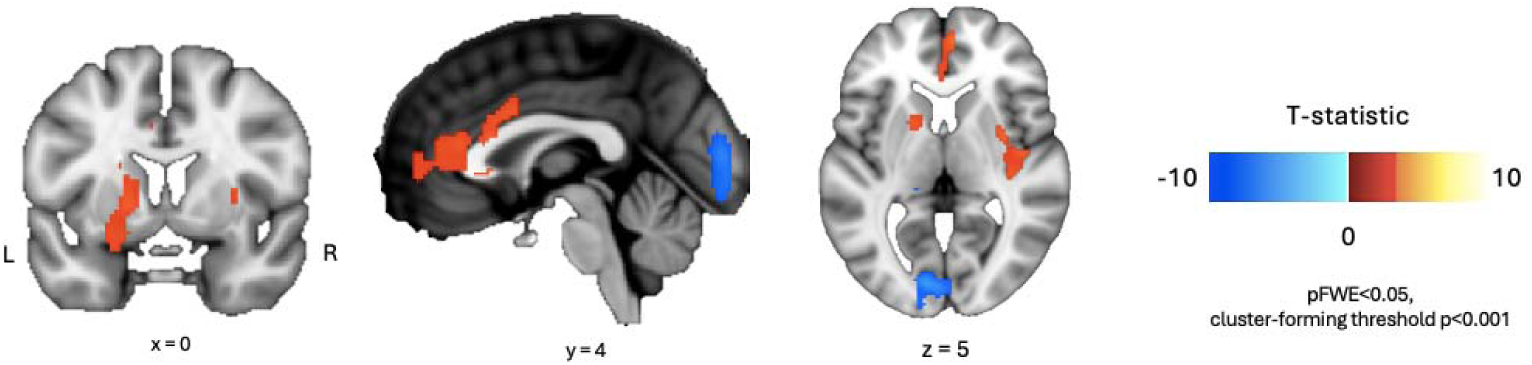
Whole-brain exploratory analysis of group differences of regional CBF. Voxelwise analysis of resting CBF revealed several significant clusters where individuals with PCS exhibited higher perfusion relative to controls. These included the anterior and mid-cingulate cortex, bilateral insula, superior temporal gyrus, precentral gyrus, right frontal cortex, thalamus, caudate, and putamen. Reduced perfusion in PCS was observed in the left posterior cingulate, occipital cortex, cerebellum, and parahippocampal gyrus. No significant group differences were observed for OEF or CMROLJ at the whole-brain corrected threshold. Clusters were considered significant for a cluster forming threshold of p<0.001 and pFWE<0.05 after multiple testing correction.

### Sensitivity Analysis on grey matter volume

No significant group effects on GMV were observed between patients and controls when accounting for age, gender, dexterity, and TIV. This suggests that the observed neurovascular and metabolic alterations in PCS are not attributable to structural atrophy.

### Associations with Symptoms, Cognition, and Blood Biomarkers

To explore the relationship between regional oxygen metabolism and behavioral, cognitive, and blood-based measures in PCS, we computed partial Spearman correlations between imaging-derived metrics and clinical variables, controlling for age, sex, handedness, and group. We focused on OEF and CMRO□ across the whole hippocampus, hippocampal subregions (whole hippocampus, CA, DG, Subiculum) and the ACC, since these were the regions for which we found the largest group differences. Because of the exploratory nature of these analyses in our pilot sample, we present the results uncorrected for multiple testing. Nevertheless, it is worth noting that none of the following reported associations would have survived strict control for multiple testing and should be thus interpreted as preliminary. Amongst variables related to clinical symptoms, significant positive correlations were observed between hippocampal DG CMRO□ and OEF values and depressive symptom severity. Additionally, CMRO□ in the ACC was negatively associated with scores of reduced motivation from the multimodal fatigue scale and with depressive symptomatology. Amongst variables related to blood analyses, CMRO_2_ in the ACC was negatively correlated with serum TNF-alpha, GFAP concentrations and erythrocyte sedimentation rate. We also found negative correlations between CMRO_2_ in the ACC and median platelet volume. CMRO_2_ in the hippocampal subiculum subregion was positively correlated with free T4 concentrations and with the monocyte-to-leucocyte ratio. Whole-hippocampus OEF and CMRO_2_ were positively correlated with the AST concentrations. Similar significant positive associations were also found for OEF in the hippocampal subfields CA and subiculum. Amongst cognition-related variables, CMRO□ in the hippocampal subiculum and ACC were positively correlated with performance on delayed memory Cognitron task. No other associations met the significance threshold of p_unc_ ≤ 0.05.

**Figure 5.**
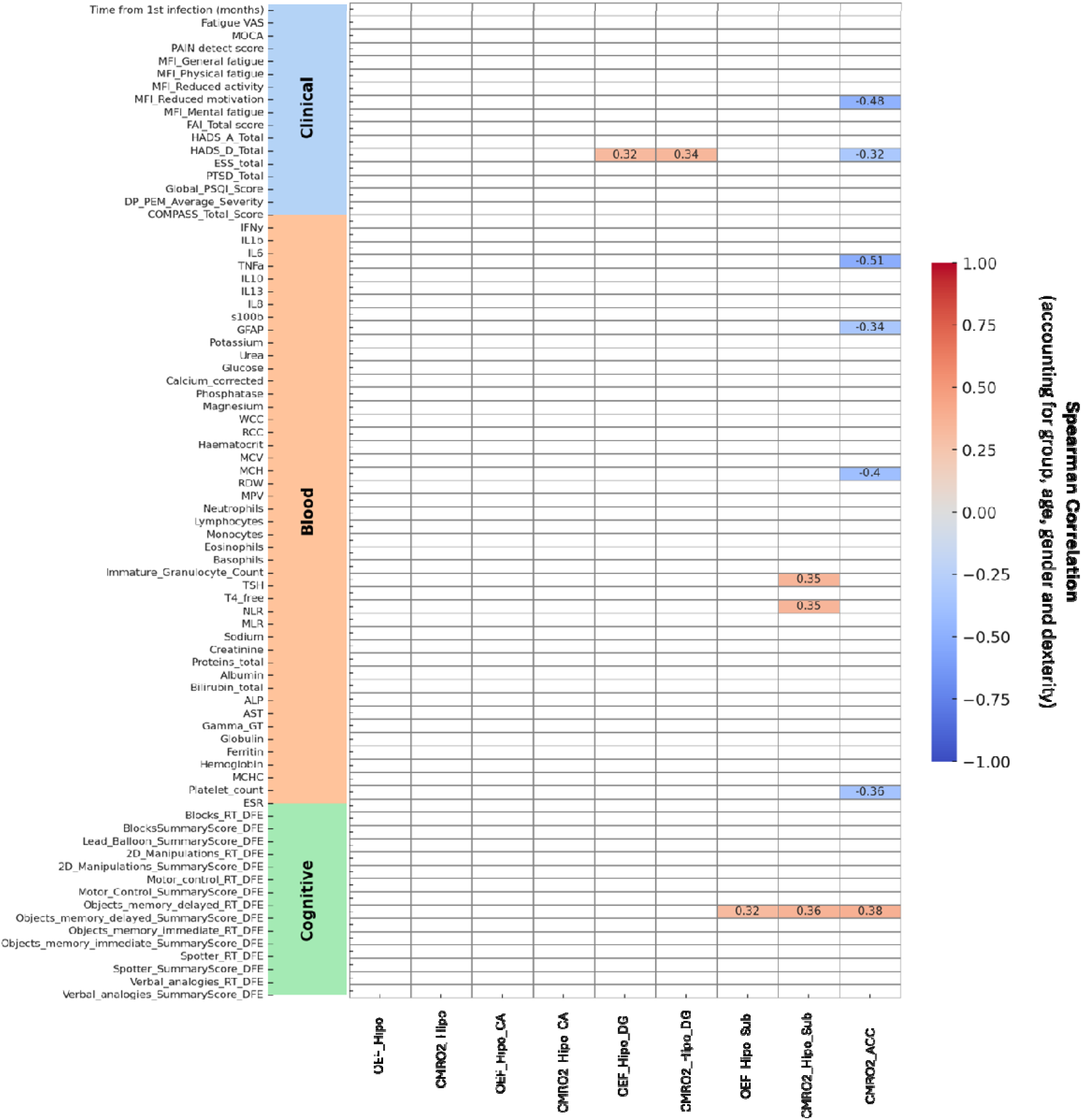
Heatmap of partial Spearman correlations between regional brain oxygen metabolism and clinical, blood-based, and cognitive variables in individuals with Post-COVID-19 Syndrome (PCS) and healthy recovered controls. The analysis includes imaging-derived metrics - oxygen extraction fraction (OEF) and cerebral metabolic rate of oxygen consumption (CMROL) - from the hippocampus (whole and subregions: CA, DG, Subiculum) and anterior cingulate cortex (ACC). Correlations were adjusted for group, age, sex, and handedness. Only statistically significant associations (p ≤ 0.05) are shown, with corresponding ρ values annotated. Non-significant cells are masked in white. Coloured bars on the left denote domain: Clinical (blue), Blood (orange), and Cognitive (green).

In addition to region-specific metabolic analyses, we explored the associations between resting cerebral blood flow (rCBF) clusters identified in the whole-brain voxelwise analysis and clinical, cognitive, and blood-based variables (Supplementary Figure S1). Cluster 1 (anterior/mid cingulate) showed positive correlations with disease duration, blood urea, free T4 levels and cognitive performance during the Spotter task, while Cluster 2 (right insula and superior temporal cortex) was positively associated with disease duration, autonomic symptoms, phosphatase level and negatively correlated with the counts of basophils and the amount of blood proteins (including albumin). Cluster 3 (left insula, thalamus, caudate) showed positive correlations with MOCA scores, MCV and MCH, and negative correlations with serum IL-10, platelet count and total proteins. Cluster 4 (right frontal operculum and putamen) only showed negative correlations with the severity of PEM. Cluster 5 (posterior cingulate, occipital, cerebellum), where PCS patients showed hypoperfusion, was negatively associated with anxiety symptoms and cognitive performance during the blocks task. Finally, Cluster 6 (hippocampal-striatal) showed a positive correlation with cognitive performance during the blocks task and a negative correlation with autonomic symptoms. No other associations met the significance threshold of p_unc_ ≤ 0.05.

## Discussion

This study used non-invasive multimodal MRI to examine resting cerebral perfusion and oxygen metabolism in individuals with PCS compared to recovered controls. While global physiological metrics were preserved, region-specific analyses revealed focal alterations in brain energetics and perfusion, particularly within circuits implicated in memory, motivation, and interoception. In a normal homeostatic regime, rCBF and OEF would compensate each other to maintain CMRO_2_, but our data shows that in some regions this is not the case. Several regions demonstrated dissociations between perfusion and metabolic demand - consistent with neurovascular uncoupling - and perfusion patterns correlated with symptom severity, immune markers, and blood rheological parameters. Together, these findings suggest that immune–vascular interactions may contribute to regional brain dysfunction in PCS, offering new targets for further research into patient stratification and metabolism-oriented therapeutic interventions.

Within the hippocampus, PCS participants exhibited significantly elevated CMRO□ and OEF across multiple subregions - including the cornu ammonis, dentate gyrus, and subiculum - in the absence of CBF changes. This pattern suggests that increased oxygen demand was met through enhanced extraction rather than increased supply, potentially reflecting compensatory upregulation of cellular metabolism or altered neuroglial coupling[49]. Given the hippocampus’s role in episodic memory and affective regulation, this adaptation may serve to maintain function under chronic physiological stress[50, 51]. Supporting this interpretation, hippocampal CMRO□ positively correlated with memory performance. However, the association between dentate gyrus metabolism and depressive symptoms raises the possibility that such compensation may carry cognitive or emotional costs. Future intervention studies are warranted to test whether strategies aimed at supporting hippocampal energy metabolism - such as targeted nutritional supplementation, metabolic enhancers, or neuromodulatory approaches - can mitigate depressive symptoms in individuals with PCS or other post-viral fatigue syndromes alike[52, 53].

In contrast, the ACC exhibited reduced CMRO□ in PCS despite increased perfusion - a hallmark of disrupted neurovascular coupling[54]. Under normal conditions, CBF is tightly linked to metabolic demand; thus, this mismatch may reflect impaired autoregulation, possibly driven by inflammatory signalling or autonomic dysregulation, which are prevalent in PCS[55, 56]. This interpretation is supported by negative correlations between ACC metabolism and inflammatory markers (TNF-α, ESR), glial activation (GFAP), and motivational and cognitive performance. Mechanistically, reduced oxygen consumption in the ACC may reflect synaptic hypoactivity, mitochondrial dysfunction, or glial metabolic imbalance - pathways increasingly implicated in neuroinflammatory conditions[57, 58]. These findings identify the ACC as a metabolically vulnerable hub in PCS. Given its central role in mood regulation and its well-established involvement in depression - particularly in inflammation-related subtypes - our results may also help explain the high burden of depressive symptoms observed in PCS and reinforce the need for metabolism-informed therapeutic strategies[59–61].

Beyond these primary regions, whole-brain voxelwise analysis identified hyperperfusion in the insula, thalamus, caudate, superior temporal gyrus, and right prefrontal cortex - areas associated with salience, interoception, and attentional control[62]. Hypoperfusion was observed in posterior cortical, cerebellar, and hippocampal-striatal regions involved in visuospatial and motor processing. These perfusion alterations were not accompanied by parallel changes in CMRO□ or OEF, suggesting redistribution of vascular resources rather than localized changes in demand. Correlations with platelet indices, plasma proteins, and IL-10 further implicate systemic immune and rheological factors in regional vascular dysregulation. However, one must note that since our data were acquired at rest, any inferences with behaviour and cognition remain inherently indirect and should be taken cautiously.

Notably, discrepancies between region-of-interest and whole-brain findings may reflect methodological differences. ROI-based analyses used bilateral anatomical masks, which may have obscured lateralized effects - such as the right-lateralized insula hyperperfusion identified in voxelwise analyses. Similarly, perfusion increases in the left ACC may have been masked by averaging across hemispheres. These findings highlight the value of integrating anatomically constrained, hypothesis-driven approaches with exploratory whole-brain analyses in studying subtle and heterogeneous effects in PCS.

Taken together, our results suggest that PCS might involve focal disruptions in brain energetics and neurovascular coupling, rather than global physiological failure. Hippocampal hypermetabolism may reflect compensatory processes that support cognitive resilience, whereas ACC dysfunction might arise from immune-glial dysregulation, contributing to motivational and affective symptoms[23]. Importantly, these abnormalities occurred in the absence of regional atrophy (as we could not detect any group-related changes in GMV), suggesting they are mostly functional in nature and might be potentially reversible (though here we are mindful that we have only assessed one single macroscopical aspect typically linked to neurodegeneration, GMV, and other microstructural metrics could have provided a more fine grained assessment). This raises the need for longitudinal studies to determine the temporal stability or resolution of these physiological alterations, and to test whether they normalize with symptom recovery or targeted intervention. Given growing evidence that Long COVID may follow a dynamic and potentially heterogeneous trajectory - characterized by partial recovery in some individuals and symptom persistence or fluctuation in others - our cross-sectional findings capture only a snapshot within this broader temporal evolution. It remains unclear whether the observed hippocampal hypermetabolism and anterior cingulate neurovascular uncoupling represent transient compensatory adaptations, longer-term maladaptive processes, or markers of recovery versus chronicity. Longitudinal studies spanning earlier and later stages of PCS, and incorporating repeated physiological imaging alongside symptom tracking, will be essential to determine the stability, progression, or reversibility of these alterations and to establish their prognostic significance. Such work would help validate hippocampal CMRO□ as a marker of compensatory capacity and ACC perfusion–metabolism mismatch as a vulnerability index. Ultimately, imaging-based biomarkers of regional metabolism may inform patient stratification, guide personalized therapeutic strategies, and enable monitoring of treatment response in future clinical trials.

This study has several limitations. First, these findings cannot be considered as specific alterations linked to PCS as we have not included a second post-viral disease control group. While the affected brain circuits are not unique to PCS, their emergence following a defined viral insult and their association with immune–vascular markers suggest PCS may represent a post-infectious instantiation of a broader, transdiagnostic neurobiological phenotype. Second, the modest sample size restricts the statistical power to detect small or spatially diffuse effects, and limits the generalizability of findings. Third, the quantification of oxygen extraction fraction (OEF) using asymmetric spin echo (ASE) methods is known to exhibit relatively low signal-to-noise ratio, which may influence the precision of CMRO□ estimates and the interpretation of regional neurovascular uncoupling. Our implementation differed from the approach proposed by Stone et al.[39] in two important respects: we employed gradient shimming along the Z-direction rather than signal averaging across thin axial slices, and we did not apply a fluid-attenuated inversion recovery (FLAIR) module to suppress cerebrospinal fluid signal. While these choices reduced acquisition time and are unlikely to introduce major artefacts, we cannot exclude the possibility that uncorrected field gradients along the X and Y axes or partial volume effects may have introduced subtle bias in OEF estimation. Fourd, physiological MRI offers a promising complementary approach to PET by enabling the simultaneous estimation of CBF, OEF, and CMRO[. However, unlike FDG-PET, which can reveal individual-level focal hypometabolism, qBOLD-derived CMRO□ and ASL-CBF remain primarily sensitive at the group level. This distinction reflects current technical constraints - including SNR limitations and model-based assumptions - that are important to consider when interpreting physiological MRI findings. Nevertheless, this scenario may evolve with the development of large-scale collaborative datasets and normative modelling frameworks, which have the potential to substantially improve single-subject precision and move physiological MRI toward more personalised inference[63–65]. Fifth, the cross-sectional design precludes causal inference and does not allow assessment of temporal dynamics in symptoms or physiological recovery. Longitudinal studies will be needed to determine whether the observed metabolic and perfusion abnormalities resolve over time, persist, or predict clinical trajectories. Sixth, exploratory associations between imaging metrics and cognitive, clinical, and peripheral measures were not corrected for multiple comparisons and should be considered hypothesis-generating. In particular, although our extraction of rCBF from voxelwise clusters avoids circularity in a strict statistical sense (i.e., independent variables were not used to define the clusters), it still introduces non-independence that may inflate correlation coefficients, as highlighted in prior literature[66]. Seventh, group differences in educational attainment must also be acknowledged: although both groups fell within the high range of educational achievement, controls exhibited significantly higher levels on average. We chose not to include education as a covariate in primary analyses due to its collinearity with group status, which poses a risk of overcorrecting and obscuring meaningful group-related effects. Importantly, none of the cognitive or perfusion/metabolism metrics correlated significantly with years of education, suggesting that observed group differences are unlikely to be driven by this variable. Finally, several potentially relevant biological and environmental factors were not systematically controlled, including viral strain, prior vaccination, use of antihistamines, hormonal status, and nutritional supplementation. These could influence immune responses, autonomic tone, or cerebral metabolism. Future research should involve larger, demographically diverse, and longitudinally followed cohorts, with comprehensive profiling of immune, endocrine, metabolic, and autonomic parameters. Multimodal imaging that integrates physiological MRI with glial PET, expanded cytokine and metabolite panels, and wearable autonomic monitoring may be especially informative for clarifying the immune–vascular–metabolic interface in PCS.

In summary, this study demonstrates that PCS is associated with regionally specific alterations in cerebral perfusion and oxygen metabolism. These findings support a dual-process model in which hippocampal hypermetabolism seems to reflect compensatory adaptation, while ACC neurovascular uncoupling might derive from neuroimmune disruption. Non-invasive physiological imaging offers a promising tool for detecting subclinical brain dysfunction in PCS and for informing personalized therapeutic strategies supporting brain metabolism.

## Credit authorship statement

**Daniel Martins**: Conceptualization, Methodology, Formal analysis, Investigation, Writing – original draft, Supervision, Project administration. **Matthew Burrows**: Methodology, Writing – review & editing. **Owen O’Daily**: Methodology, Writing – review & editing. **Ziyuan Cai**: Data processing, Quality control, Writing – review & editing. **Nicole Mariani**: Methodological support, Writing – review & editing. **Alessandra Borsini**: Methodological support, Writing – review & editing. **Valeria Mondelli**: Conceptualization, Writing – review & editing. **Brandi Eiff**: Participant recruitment, Investigation, Data curation. **Silvia Rota**: Participant recruitment, Clinical assessments, Project coordination. **Timothy Nicholson**: Clinical oversight, Participant recruitment, Writing – review & editing. **Laila Rida**: Methodology, Writing – review & editing. **Adam Hampshire**: Cognitive task development, Normative modeling, Software, Writing – review & editing. **Federico Turkheimer**: Statistical supervision, Interpretation, Writing – review & editing. **Catherine Morgan:** Methodology, Writing – review & editing. **David Lythgoe**: Methodology, Writing – review & editing. **Steven C. R. Williams**: Resources, Funding acquisition, Supervision, Writing – review & editing. **Fernando Zelaya**: Conceptualization, Methodology, Supervision, Writing – review & editing. All authors have read and approved the final manuscript and agree to be accountable for all aspects of the work.

## Acknowledgements

This study was funded by the National Institute for Health and Care Research (NIHR) Maudsley Biomedical Research Centre (BRC), South London and Maudsley NHS Foundation Trust, under the “Reach Out” funding call (Ref: R0-01). We are grateful to all participants for their time and commitment to the study. We thank the radiographers and technical staff at the Centre for Neuroimaging Sciences (King’s College London) for their assistance with MRI acquisition. We also acknowledge Synnovis (Viapath) for processing routine blood analyses, the Clinical Research Facility (CRF) team at King’s College Hospital Foundation Trust for their support during participant visits and blood collection, and the Cognitron platform team for assistance with cognitive data management and normative modeling. Dr Alessandra Borsini is funded by the UK Medical Research Council (grants MR/L014815/1, MR/J002739/1 and MR/N029488/1), the European Commission Horizon 2020 (grant SC1-BHC-01-2019), the Psychiatric Research Trust (grant RE23747), the Rosetrees Trust (grants CF-2023-I-2\101, Seedcorn2024\100004) and the National Institute for Health Research (NIHR) Biomedical Research Centre at South London and Maudsley NHS Foundation Trust and King’s College London.

