## Supplementary for "Multimodal imaging suggests potential immune-vascular contributions to altered regional brain perfusion and oxygen metabolism in Post-COVID-19 Syndrome"

**Supplementary Table S1. Self-reported fatigue, affective symptoms, and somatic complaints in PCS and HC participants, with Bayesian inference.** Comparison of symptom severity between individuals with Post-COVID-19 Syndrome (PCS) and matched healthy controls (HC) across domains including fatigue (VAS, MFI), sleep (PSQI), autonomic function (COMPASS-31), post-exertional malaise, mood (HADS), PTSD, and fibromyalgia-related symptoms. Values are shown as mean (standard deviation), and group comparisons were conducted using independent samples t-tests or chi-squared tests, as appropriate. The Bayesian factor (BF₁₀) is reported for each comparison, quantifying the strength of evidence in favor of the alternative hypothesis. Asterisks (*) denote statistically significant differences at p < 0.05 or BF₁₀ > 1.

| **Variable** | **PCS**  **Mean (std)** | **HC**  **Mean (std)** | **Statistics** | **P-value** | **B_10_** |
| --- | --- | --- | --- | --- | --- |
| **Fatigue VAS** | 5.500 (1.606) | 1.850 (1.663) | T(38)=7.061 | 2.036x10^-8^* | 876.874* |
| **Multidimensional fatigue**  **General** **Physical**  **Reduced activity**  **Reduced motivation**  **Mental fatigue** | 15.200 (3.205)  14.600 (2.998)  12.150 (3.703)  8.600 (3.393)  12.850 (3.843) | 6.684 (2.212)  5.737 (2.884)  5.444 (2.307)  4.895 (2.378)  6.053 (3.027) | T(38)=47.68  T(38)=43.19  T(38)=15.09  T(38)=9.59  T(38)=21.41 | 1.355x10^-11*^  2.408x10^-11^*  1.072x10^-7*^  3.589x10^-4^*  4.391x10^-7^* | 1332.772*  4158.129*  650.746*  34.338*  767.860* |
| **ESS** | 12.400 (5.548) | 6.474 (4.234) | T(37)=3.735 | 6.304x10^-4^* | 24.874* |
| **HADS Depression** | 8.900 (2.954) | 4.053 (2.345) | T(37)=5.657 | 1.833x10^-6^* | 153.950* |
| **HADS Anxiety** | 7.550 (3.790) | 5.368 (2.477) | T(37)=2.116 | 0.041* | 1.287* |
| **PTSD total score** | 19.750 (10.208) | 6.105 (8.185) | T(37)=4.590 | 4.964x10^-5*^ | 59.715* |
| **De Paul's**  **PEM** | 2.950 (0.945) | 0.421 (0.449) | T(37)=10.584 | 9.541x10^-13*^ | 64546.865* |
| **De Paul's**  **Unrefreshing sleep** | 3.400 (0.821) | 1.316 (0.749) | T(37)=8.269 | 6.229x10^-10*^ | 1547.301* |
| **De Paul's**  **Autonomic symptoms** | 1.775 (0.881) | 0.553 (0.550) | T(37)=5.166 | 8.438x10^-6*^ | 73.625* |
| **De Paul's**  **Neurocognitive** | 1.775 (0.910) | 0.474 (0.612) | T(37)=5.212 | 7.316x10^-6*^ | 401.197* |
| **De Paul's**  **Immune** | 2.250 (0.993) | 0.658 (0.834) | T(37)=5.405 | 4.010x10^-6*^ | 381.905* |
| **COMPASS total score** | 62.413 (19.629) | 30.666 (19.158) | T(37)=5.108 | 1.011x10^-5*^ | 569.590* |
| **Global PSQI** | 15.200 (6.542) | 9.263 (5.791) | T(37)=2.995 | 0.005* | 15.254* |
| **MRC**  **Dyspnea**  **Scale** | Stage 1: 9  Stage 2: 9  Stage 3: 2 | 19  1  0 | χ^2^(5)=11.971 | 0.003* | N/A |
| **MoCA** | 26.8 (2.55) | 26.5 (1.54) | T(37)=0.430 | 0.668 | 0.420 |
| **Pain detect**  **WPI**  **SS** | 9.00(6.53)  5.45 (4.07)  8.55 (1.88) | 1.7 (1.84)  1.00 (1.62)  1.75 (1.68) | T(37)= 4.813  T(37)=4.541  T(37)=12.066 | <0.001*  <0.001*  <0.001* | 1315.956*  119.442*  13663.606* |
| **Fibromyalgia**  **diagnostic criteria** | Yes: 8  No: 12 | Yes:0  No:20 | χ^2^(2)=10.581 | 0.005* | N/A |

**Supplementary Table S2. Comparison of standard blood parameters between PCS and healthy control participants.** Descriptive statistics and group comparisons for hematological, biochemical, and inflammatory markers in individuals with Post-COVID-19 Syndrome (PCS) and matched healthy controls (HC). Results are reported as mean (standard deviation). Between-group comparisons were conducted using the Mann–Whitney U test (W), with corresponding p-values and Bayesian factors (BF₁₀) reported for each variable. Asterisks (*) denote statistically significant differences at p < 0.05 or BF₁₀ > 1.

| **Variable** | **PCS Mean (SD)** | **HC Mean (SD)** | **Statistics**  **W** | **P-value** | **B_10_** |
| --- | --- | --- | --- | --- | --- |
| **Sodium** | 140.118 (2.619) | 140.474 (2.091) | 161.00 | 1.00 | 0.369 |
| **Potassium** | 3.965 (0.237) | 4.000 (0.252) | 126.00 | 0.53 | 0.334 |
| **Urea** | 4.135 (1.346) | 4.337 (0.942) | 158.50 | 0.94 | 0.359 |
| **Creatinine** | 61.235 (8.318) | 68.632 (11.847) | 101.50 | 0.06 | 1.232* |
| **Glucose** | 4.871 (0.819) | 4.968 (1.106) | 157.00 | 0.90 | 0.329 |
| **Calcium_corrected** | 2.268 (0.059) | 2.288 (0.109) | 159.00 | 0.95 | 0.341 |
| **Phosphatase** | 1.088 (0.166) | 1.171 (0.163) | 106.00 | 0.08 | 0.749 |
| **Proteins_total** | 71.000 (3.691) | 71.526 (3.878) | 148.50 | 0.69 | 0.337 |
| **Albumin** | 47.000 (1.969) | 47.000 (2.749) | 154.50 | 0.83 | 0.326 |
| **Bilirubin_total** | 10.125 (6.672) | 9.824 (7.170) | 116.00 | 0.48 | 0.370 |
| **ALP** | 64.176 (15.204) | 63.947 (23.710) | 145.50 | 0.62 | 0.362 |
| **AST** | 23.353 (6.441) | 22.316 (7.521) | 131.00 | 0.34 | 0.420 |
| **Gamma_GT** | 13.059 (3.508) | 31.632 (47.978) | 97.50 | 0.04 | 1.735* |
| **Magnesium** | 0.859 (0.043) | 0.872 (0.054) | 151.50 | 0.76 | 0.347 |
| **CRP_recoded** | 1.353 (0.606) | 1.368 (0.597) | 158.50 | 0.92 | 0.397 |
| **Globulin** | 23.529 (3.875) | 24.526 (2.674) | 138.50 | 0.47 | 0.497 |
| **Ferritin** | 108.353 (103.511) | 99.789 (84.020) | 156.50 | 0.89 | 0.307 |
| **WCC** | 6.629 (1.662) | 6.501 (1.803) | 156.50 | 0.89 | 0.336 |
| **RCC** | 4.385 (0.457) | 4.561 (0.386) | 112.50 | 0.12 | 0.843 |
| **Hemoglobin** | 134.000 (13.224) | 137.263 (10.514) | 138.50 | 0.48 | 0.426 |
| **Haematocrit** | 0.403 (0.036) | 0.417 (0.029) | 120.00 | 0.19 | 0.739 |
| **MCV** | 91.988 (4.276) | 91.679 (5.684) | 159.00 | 0.95 | 0.329 |
| **MCH** | 30.624 (1.719) | 30.200 (2.149) | 151.00 | 0.75 | 0.348 |
| **MCHC** | 332.882 (11.931) | 329.158 (8.662) | 128.50 | 0.30 | 0.504 |
| **RDW** | 12.406 (0.599) | 12.474 (0.726) | 154.50 | 0.84 | 0.329 |
| **Platelet_count** | 267.118 (42.355) | 289.789 (45.281) | 105.00 | 0.08 | 1.123* |
| **MPV** | 10.988 (0.834) | 10.716 (0.727) | 121.00 | 0.20 | 0.463 |
| **Neutrophils** | 4.142 (1.178) | 3.773 (1.409) | 135.50 | 0.42 | 0.414 |
| **Lymphocytes** | 1.934 (0.550) | 2.029 (0.650) | 144.50 | 0.60 | 0.334 |
| **Monocytes** | 0.441 (0.102) | 0.441 (0.127) | 155.00 | 0.85 | 0.319 |
| **Eosinophils** | 0.141 (0.100) | 0.214 (0.243) | 147.50 | 0.67 | 0.365 |
| **Basophils** | 0.052 (0.030) | 0.042 (0.018) | 136.00 | 0.42 | 0.486 |
| **Immature_Granulocyte_Count** | 0.022 (0.016) | 0.022 (0.012) | 141.50 | 0.51 | 0.327 |
| **ESR** | 7.588 (6.662) | 8.278 (8.072) | 139.50 | 0.67 | 0.354 |
| **TSH** | 1.819 (0.630) | 1.633 (0.658) | 124.50 | 0.25 | 0.423 |
| **T4_free** | 15.124 (1.759) | 15.042 (1.624) | 156.00 | 0.87 | 0.321 |
| **NLR** | 2.209 (0.594) | 2.038 (1.137) | 125.50 | 0.26 | 0.540 |
| **MLR** | 0.234 (0.041) | 0.230 (0.072) | 156.00 | 0.87 | 0.346 |

**Supplementary Table S3. Serum cytokines and glial biomarkers in PCS and healthy control participants.** Group comparisons for circulating immune markers (interleukins, interferon-γ, TNF-α) and glial markers (S100β, GFAP) measured in serum samples from individuals with Post-COVID-19 Syndrome (PCS) and healthy controls (HC). Data are reported as mean (standard deviation). Concentrations are in pg/mL. Between-group differences were evaluated using the Mann–Whitney U test (W), with corresponding p-values and Bayes Factors (BF₁₀). Asterisks (*) indicate statistically suggestive effects (BF₁₀ > 1)

| **Variable** | **PCS Mean (SD)** | **HC Mean (SD)** | **Statistics**  **W** | **P-value** | **B_10_** |
| --- | --- | --- | --- | --- | --- |
| **IFNy** | 4.639 (8.032) | 4.182 (8.177) | 151.000 | 0.789 | 0.345 |
| **IL1b** | 0.345 (0.745) | 0.094 (0.061) | 195.000 | 0.300 | 0.573 |
| **IL6** | 1.197 (2.795) | 0.418 (0.205) | 194.000 | 0.478 | 0.431 |
| **TNFa** | 1.164 (1.799) | 0.980 (0.316) | 110.000 | 0.116 | 1.094* |
| **IL10** | 0.576 (1.145) | 0.229 (0.141) | 171.000 | 0.449 | 0.365 |
| **IL13** | 0.998 (0.889) | 1.228 (0.894) | 127.000 | 0.563 | 0.398 |
| **IL8** | 6.106 (4.671) | 6.581 (2.791) | 126.000 | 0.289 | 0.460 |
| **s100b** | 3.908 (2.192) | 5.165 (2.558) | 130.000 | 0.149 | 1.025* |
| **GFAP** | 14.62 (4.79) | 13.78 (5.32) | 157.000 | 0.515 | 0.406 |

**Supplementary Table S4. Performance on computerized cognitive tasks in PCS and healthy control participants.** Comparison of standardized deviation-from-expected (DFE) scores derived from the Cognitron cognitive task battery, assessing domains such as spatial reasoning, memory, motor control, and sustained attention. Scores represent z-standardized residuals relative to normative models controlling for age, sex, handedness, and ethnicity. Between-group differences were tested using independent samples t-tests, with corresponding p-values and Bayes Factors (BF₁₀). Asterisks (*) indicate results meeting the threshold for statistical significance (p < 0.05) or annedoctal Bayesian evidence (BF₁₀ > 1).

| **Variable** | **PCS Mean (SD)** | **HC Mean (SD)** | **Statistics** | **P-value** | **B_10_** |
| --- | --- | --- | --- | --- | --- |
| **Blocks_RT_DFE** | 0.136 (0.613) | -0.043 (0.539) | T(38)=0.981 | 0.33 | 0.452 |
| **BlocksSummaryScore_DFE** | -0.706 (1.069) | -0.301 (1.011) | T(38)=-1.231 | 0.23 | 0.563 |
| **Lead_Balloon_SummaryScore_DFE** | 0.576 (1.254) | -0.038 (0.658) | T(38)=1.939 | 0.06 | 1.330* |
| **2D_Manipulations_RT_DFE** | -0.006 (0.741) | 0.599 (1.892) | T(38)=-1.332 | 0.20 | 0.621 |
| **2D_Manipulations_SummaryScore_DFE** | 1.573 (1.484) | 1.236 (1.711) | T(38)=0.665 | 0.51 | 0.369 |
| **Motor_control_RT_DFE** | -0.097 (0.740) | -0.148 (0.766) | T(38)=0.214 | 0.83 | 0.314 |
| **Motor_Control_SummaryScore_DFE** | -0.012 (0.772) | -0.122 (0.764) | T(38)=0.453 | 0.65 | 0.335 |
| **Objects_memory_delayed_RT_DFE** | 0.416 (1.384) | -0.458 (0.898) | T(38)=2.369 | 0.02* | 2.653* |
| **Objects_memory_delayed_SummaryScore_DFE** | 0.073 (1.025) | -0.386 (1.866) | T(38)=0.964 | 0.34 | 0.446 |
| **Objects_memory_immediate_RT_DFE** | -0.106 (0.757) | -0.433 (0.840) | T(38)=1.293 | 0.20 | 0.596 |
| **Objects_memory_immediate_SummaryScore_DFE** | 0.023 (0.845) | -0.006 (1.268) | T(38)=0.085 | 0.93 | 0.310 |
| **Spotter_RT_DFE** | 0.360 (1.302) | -0.308 (0.841) | T(38)=1.927 | 0.06 | 1.303* |
| **Spotter_SummaryScore_DFE** | 0.093 (0.175) | 0.120 (0.361) | T(38)=-0.301 | 0.77 | 0.320 |
| **Verbal_analogies_RT_DFE** | 0.116 (0.743) | -0.027 (0.916) | T(38)=0.542 | 0.59 | 0.347 |
| **Verbal_analogies_SummaryScore_DFE** | -0.852 (1.355) | -0.592 (1.163) | T(38)=-0.651 | 0.52 | 0.366 |

**Supplementary Table S5. Whole-brain voxelwise group differences in regional cerebral blood flow (CBF) between PCS and healthy control participants.** Significant clusters identified in whole-brain voxelwise analysis comparing CBF between Post-COVID-19 Syndrome (PCS) participants and matched healthy controls (HC), accouting for age, gender, dexterity and global CBF. Coordinates represent peak voxels in MNI space. Cluster-level family-wise error (pFWE) correction was applied at p < 0.05, with a cluster-forming threshold of p<0.001.

| **Contrast** | **Region** | **Cluster label** | **pFWE** | **k** | **Z** | **X**  **mm** | **Y**  **mm** | **Z**  **mm** |
| --- | --- | --- | --- | --- | --- | --- | --- | --- |
| **PCS > HC** | Middle Frontal gyrus, Anterior, Middle Cingulate | 1 | <0.001 | 881 | 4.55 | 6 | 52 | 8 |
|  |  |  |  |  |  | -4 | 10 | 34 |
|  |  |  |  |  |  | -6 | 42 | 18 |
|  | Right insula, superior temporal, precentral gyrus | 2 | <0.001 | 570 | 4.38 | 40 | -18 | -6 |
|  |  |  |  |  |  | 34 | -2 | 4 |
|  |  |  |  |  |  | 46 | -10 | 8 |
|  | Left insula, Left thalamus, left caudate | 3 | 0.004 | 373 | 3.97 | -16 | 4 | 2 |
|  |  |  |  |  |  | -20 | 6 | -14 |
|  |  |  |  |  |  | -20 | 0 | 18 |
|  | Right middle frontal and inferior gyri, insular cortex/putamen | 4 | 0.017 | 277 | 4.62 | 48 | 14 | 44 |
|  |  |  |  |  |  | 52 | 20 | 28 |
|  |  |  |  |  |  | 42 | 14 | 16 |
| **PCS < HC** | Left occipital, left posterior cingulate, left cerebellum | 5 | <0.001 | 1010 | 4.84 | -10 | -86 | 4 |
|  |  |  |  |  |  | -6 | -84 | 24 |
|  |  |  |  |  |  | -14 | -94 | 6 |
|  | Left parahippocampal gyrus, left striatum | 6 | <0.001 | 865 | 4.22 | -16 | -68 | -12 |
|  |  |  |  |  |  | -24 | -50 | -18 |
|  |  |  |  |  |  | -14 | -38 | -2 |

**Supplementary Figure S1. Associations between perfusion clusters and clinical, blood-based, and cognitive variables.** Partial Spearman correlations between voxelwise perfusion clusters (identified in whole-brain CBF analysis; see Table S5) and clinical, blood-based, and cognitive variables. Correlations were adjusted for group, age, sex, and manual dexterity. Anatomical labels for each significant perfusion cluster (Clusters 1–6) are shown on the left in MNI space. The heatmap on the right displays correlation coefficients (color-coded from –1.0 to +1.0), with only statistically significant associations (p < 0.05, uncorrected) annotated with numerical values. Domains are color-coded: clinical (blue), blood biomarkers (orange), and cognitive task performance (green).

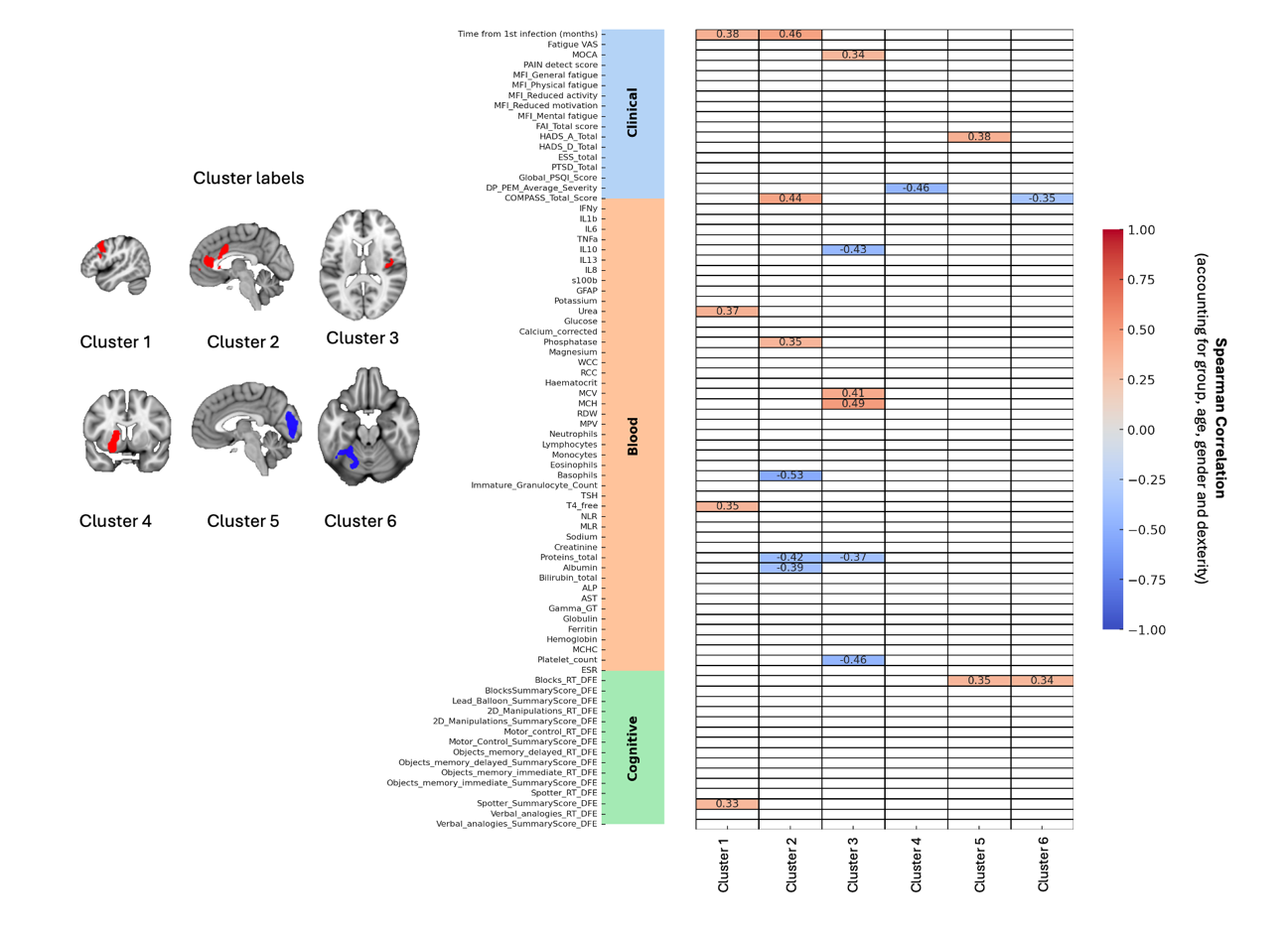
